# Native extracellular matrix probes to target patient- and tissue- specific cell-microenvironment interactions by force spectroscopy

**DOI:** 10.1101/2022.12.02.518867

**Authors:** H. Holuigue, L. Nacci, P. Di Chiaro, M. Chighizola, I. Locatelli, C. Schulte, M. Alfano, G.R. Diaferia, A. Podestà

## Abstract

Atomic Force Microscopy (AFM) is successfully used for the quantitative investigation of the cellular mechanosensing of the microenvironment. To this purpose, several force spectroscopy approaches aim at measuring the adhesive forces between two living cells and also between a cell and a suitable reproduction of the extracellular matrix (ECM), typically exploiting tips suitably functionalised with single components (e.g. collagen, fibronectin) of the ECM. However, these probes only poorly reproduce the complexity of the native cellular microenvironment and consequently of the biological interactions.

We developed a novel approach to produce AFM probes that faithfully retain the structural and biochemical complexity of the ECM; this was achieved by attaching to an AFM cantilever a micrometric slice of native decellularised ECM, which was cut by laser microdissection. We demonstrate that these probes preserve the morphological, mechanical, and chemical heterogeneity of the ECM.

Native ECM probes can be used in force spectroscopy experiments aimed at targeting cell-microenvironment interactions. Here, we demonstrate the feasibility of dissecting mechanotransductive cell-ECM interactions in the 10 pN range. As proof-of-principle, we tested a rat bladder ECM probe against the AY-27 rat bladder cancer cell line. On the one hand, we obtained reproducible results using different probes derived from the same ECM regions; on the other hand, we detected differences in the adhesion patterns of distinct bladder ECM regions, such as submucosa and detrusor, in line with the disparities in composition and biophysical properties of these ECM regions.

Our results demonstrate that native ECM probes, produced from patient-specific regions of organs and tissues, can be used to investigate cell-microenvironment interactions and early mechanotransductive processes by force spectroscopy. This opens new possibilities in the field of personalised medicine.

## 1. Introduction

Cells and their microenvironment have a strong intercommunication and interplay, influencing each other. The extracellular matrix (ECM) surrounding the cells is mainly composed of a three-dimensional network of collagen and crosslinked proteins, such as fibronectin and laminin; ECM from various tissues can have strong differences in their composition and biophysical properties (i.e., rigidity and structural features, such as nanotopography) ^1–4^. Cells use the ECM as a scaffold for their anchorage and interact with it in a reciprocal manner. Cells can convert external mechanical and topographical stimuli into biochemical signalling, which often determine changes in the mechanical properties of the cell through the reorganisation of the cytoskeleton and modulates gene/protein expression, a process called mechanotransduction. Cells are therefore able to perceive changes in the physical properties of their surrounding ECM. The cell itself uses force to sense the biophysical characteristics of the ECM. The study of these reciprocal interactions pertains to the fields of cellular mechanobiology ^5–12^.

Different techniques are used to study these cell-ECM interactions, among which Atomic Force Microscopy (AFM) stands out due to its ability to both sense and apply forces at the nanoscale, with sub-nanonewton sensitivity ^13–17^. In the context of biophysical investigations, many AFM-based force spectroscopy (FS) configurations can be found ^18^: a single-cell probe approaching a substrate coated with proteins ^19–24^; an ECM-mimicking probe approaching an adherent cell ^11,25–29^; a single-cell probe approaching an adherent cell ^15,30,31^. Both cell-cell and cell-ECM interactions can be investigated, through the direct measurement of faint (10-100 pN) interaction forces, which are mostly cadherin or integrin related, respectively ^32–36^.

The main advantage of FS techniques using tips functionalised with single proteins ^26,28,29^ is the simplification of the complex cell-microenvironment interface, allowing us to study one specific molecular interaction at a time. An example for this approach for the study of cell-ECM interactions consists in the functionalisation of AFM tips with collagen, a major component of the ECM ^29,37^. While these methods enable an accurate characterisation of specific molecular interactions, the reconstituted interface is poorly mimicking the complexity of the native interface in physiological conditions.

Aiming at reproducing the native cell-ECM interface within a typical AFM-based FS experiment, we developed native ECM probes and demonstrated that they can be reliably used to scrutinise integrin-related adhesive interactions between cells and their microenvironment. Attaching the ECM to the cantilever and ramping it against adherent living cells, instead of using single-cell functionalised cantilevers against an ECM sample, is advantageous, among other reasons, because it allows to test i) many different cells with the same probe, that can be used and re-used repeatedly, and ii) cells seeded on different substrates, i.e. polarized or with different phenotypes such as type 1 and type 2 macrophages or epithelial cancer cells or cancer cells during and after epithelial-mesenchymal transition. Indeed, the ECM is more stable and easier to handle than living cells, and each single-cell probe requires a different cantilever and a new calibration; overall, using ECM probes more likely provides improved reliability and statistics of the FS experiments.

Here we describe the fabrication approach of novel native ECM probes and the results of their characterisation, showing that the chemicophysical properties of the native ECM are preserved, also after intense and prolonged use. We also report on the application of these new probes in adhesion force spectroscopy experiments, showing that the peculiar patterns of adhesive molecular interactions observed in FS experiments were reproduced. As a proof-of-principle, we focused on rat bladder - derived ECM and used the rat bladder tumour cell line AY-27 as a suitable interaction partner. The described protocol is, however, universally applicable, and allows to reliably fabricate probes out of any ECM, and even from specific regions of the same ECM, to be tested against both immortalised and primary cells, paving the way to the investigation of patient-specific cell-ECM interactions.

## 2. Results and Discussion

### 3.1 Production of native ECM-probes for adhesion force spectroscopy

To produce native ECM probes, we developed a novel approach based on the cutting of ECM pieces from a glass-supported, decellularised ECM slice; these pieces will then be detached from the glass slide and attached to tipless functionalised cantilevers (Figure 1A-D).

**Figure 1.**
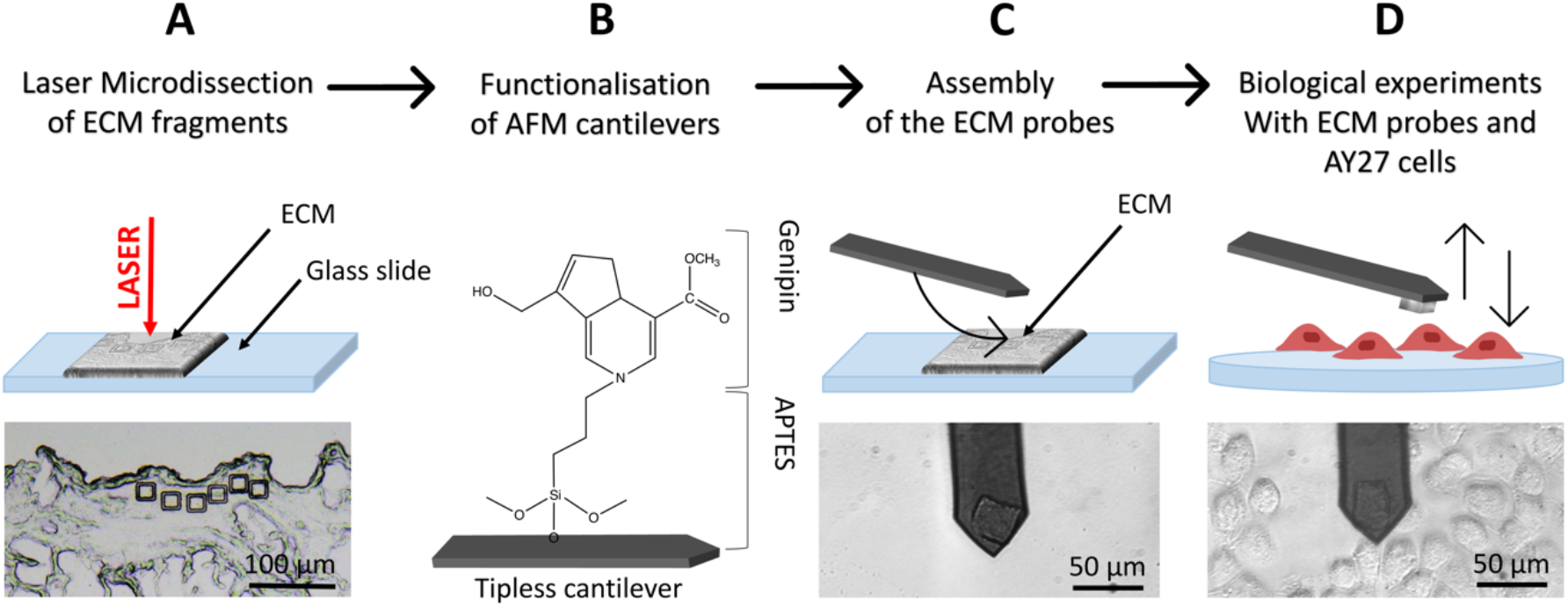
Schematics of the ECM probe production. (A) Optimised use of the LMD system to produce native ECM-probes, keeping the ECM directly on the standard microscope glass slide, used without the plastic membrane, resulting in ECM fragments physically separated from the surrounding matrix. (B) Tipless cantilevers are functionalised with APTES and genipin (covalently bonded), and then used for the attachment of the ECM probes (C), exploiting the XY micro-translation stage of the optical microscope that includes the AFM (stand-alone stages can be used equivalently). (D) The resulting ECM-probes are used for biological experiments, such as force spectroscopy measurements on top of living cells.

To cut pieces with predefined and reproducible dimensions and shapes from the ECM slice, we used an LMD apparatus (in Figure 1A, square cuts with 20μm x 20μm area are shown).

LMD is typically used to cut a selected area from a dehydrated tissue section attached on the upper surface of a glass slide, where a thin plastic membrane is cemented to the glass at its edges leaving a rectangular area with air trapped between the membrane and the glass backbone (Figure S1A-C). A focused UV laser then cuts the selected regions of the tissue, together with the plastic membrane allowing the microdissected pieces to fall by gravity in a reservoir located below. These samples are then typically digested for genomics or proteomics assays ^38,39^, where the presence of the rigid plastic membrane does not interfere with the subsequent molecular reactions.

For our application, given that the objective is to transfer and attach intact fragments of ECM to either AFM tipless cantilevers or spherical tips, the presence of the plastic membrane is detrimental. We therefore modified the standard LMD procedure to develop an optimised approach, based on the elimination of the plastic membrane from the glass slide and the introduction of suitable procedures to detach the cut pieces from the slide and to transfer them to the AFM cantilever or tip.

The protocol is represented schematically in Figure 1. Here, the laser is still used for the cut, but the ECM slices are attached directly on the glass slide, without the plastic membrane in between (Figures 1A and S2A) and placed on the microscope, with the tissue facing up. After the cut, the microdissected pieces remain on the glass surface (Figures 1A and S2B-D), separated from the surrounding ECM by a narrow empty region vaporised by the laser, where they can be picked up by, and deposited onto, the functionalised cantilever used for the assembling of the ECM probe, exploiting the XY motorised stage of the AFM integrated in the optical microscope (see Supporting Movies SM1-SM3). This procedure takes place in liquid. After the attachment, the cantilever is withdrawn from the surface and kept at rest for 10 min. ECM probes were typically stored in PBS at 37°C in a humid environment (for example in the cell incubator) and used in the following days.

The fact that the ECM pieces remain on the glass slide after the cut facilitates the investigation of same or twin specimens by other techniques, like immunofluorescence. Moreover, having the ECM attached to the cantilever and testing it against a cell population cultured on a suitable support is very convenient compared to having a single cell probe that interrogates an ECM sample; indeed, in the first case many cells can be tested with the same ECM probes, while in the standard configuration each new tested cell requires the preparation of a new probe.

In this work, 20μm x 20μm, 10μm thick squares (Figure 1A) were cut in different regions of decellularised rat bladder: submucosal, detrusor and adventitia ^17,40,41^.

### 3.2 Characterisation of ECM and ECM probes

LMD provides many advantages including the possibility to precisely select the ROI, cut ECM pieces with predefined, reproducible geometry and dimensions, and collect the samples without damaging them. To ensure that the ECM pieces retain the properties of the pristine ECM, we performed several assessments of biophysical and compositional parameters.

Firstly, we controlled the possible effect of the laser cut on morphology, mechanics and composition of the ECM, respectively, through combined AFM topographic and nanomechanical measurements (Figure 2) and quantitative immunofluorescence staining (Figure 3). For the nanomechanical assessment of the ECM, we measured the region inside and outside the microdissected piece; the inner region is the one that it is then attached to the AFM cantilever, while the outer region is taken as reference for the untreated ECM. For the immunofluorescence analysis, we measured the fluorescent signals detected after immunostaining from the inner regions of the microdissected pieces and the same regions from a consecutive section obtained from the same ECM specimen not subjected to laser ablation.

**Figure 2.**
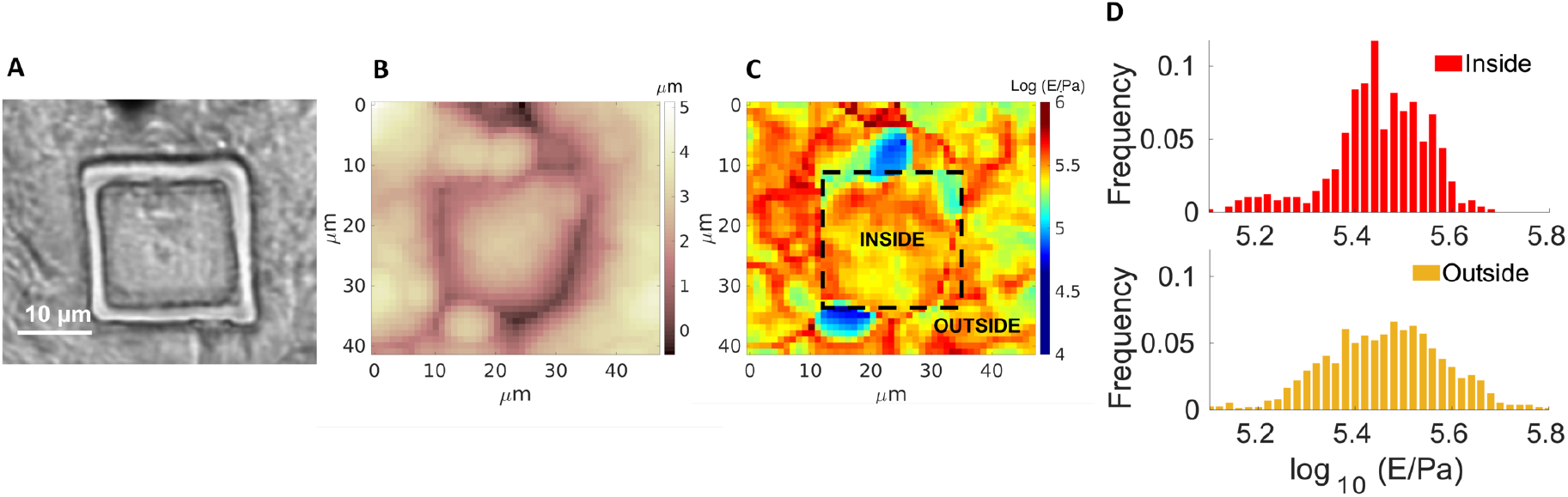
(A) Optical image, (B) Young’s modulus and (C) topographic maps of an ECM slice with the cut of the laser clearly visible (highlighted in C). (D) Distribution of the measured Young’s Modulus values for the internal and external regions of the cut.

**Figure 3.**
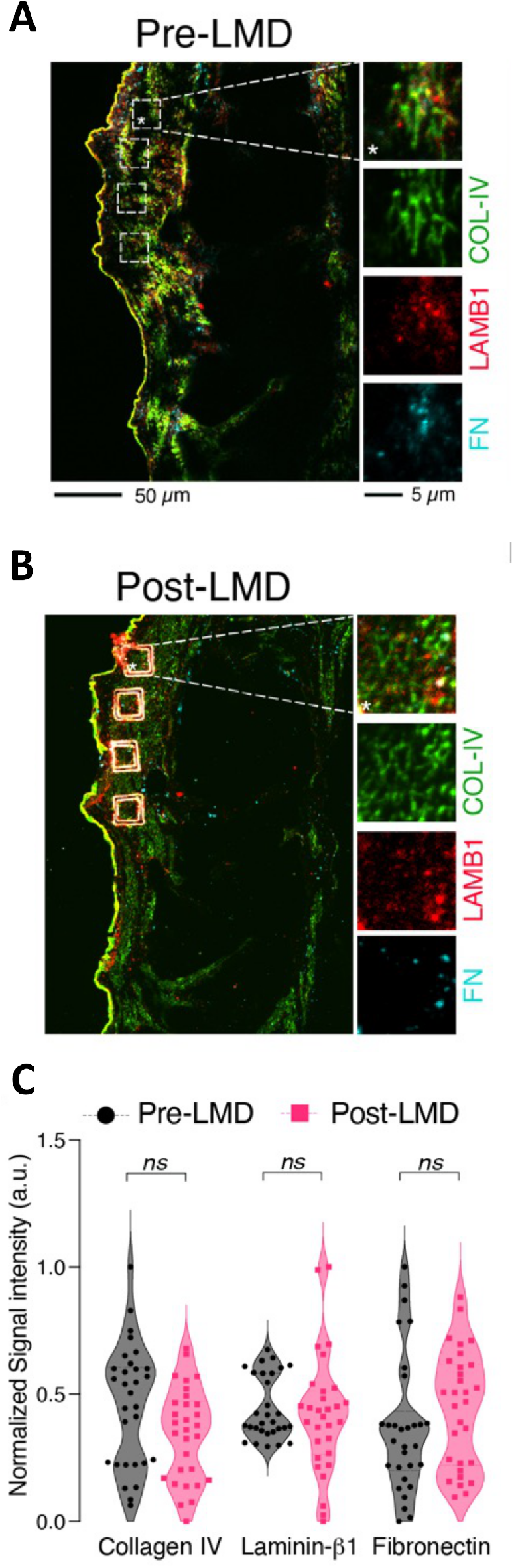
Immunostaining for Collagen IV (COL-IV, *green*), laminin β1 (LAMB1, *red*), and fibronectin (FN, *cyan*) before (A, Pre-LMD) and after (B, Post-LMD) the cut. The white boxes in the Pre-LMD image represent the matched area subjected to microdissection. (C) Quantification of the fluorescent signal associated to the different proteins in each ROI (white squares). The data represent normalised fluorescent signal (a.u., arbitrary units) scaled between 0 and 1 (n=30, 5 ROIs per each field, 6 fields acquired distributed along the layers of the bladder).

Figure 2A-C shows the optical image of a cut ECM piece together with AFM topographic and mechanical maps. The borders of the cut region are clearly visible in both optical image and topographic AFM maps. No evident differences between the inner and the outer regions are visible, regarding both morphological and mechanical properties (the median Young’s modulus values are 280 and 289 kPa for the internal and external parts, respectively). Besides the medians, also the distributions of Young’s modulus values E (Figure 2D) are similar (the distribution of the external part is slightly broader, which can be explained by the larger covered area, encompassing a larger heterogeneity of the sample).

These results are in line with the results obtained by quantitative staining of major ECM proteins such as collagen, laminin, and fibronectin on different cut pieces of ECM, demonstrating that the composition of the matrix is not affected by the cut (Figure 3). The UV laser of the LMD apparatus could induce thermal damage and autofluorescence ^42^, and fibronectin inactivation ^43^ around the edge of the ablated region; nevertheless, the central area used for the production of the probes and interacting with the cells is not affected by these phenomena.

Stress tests were performed to assess the firm attachment of the matrix to the cantilever and the force spectroscopy functionality after repeated use. The ECM remained attached to the cantilever after scanning continuously in Contact Mode on a glass slide for 1h (Figure S3A); similar adhesion forces were measured during continuous acquisition of 400 FCs (Figure S3B).

To further assess the lifetime of ECM probes, in particular the structural resistance of the ECM fragments, the firmness of their attachment to the cantilevers and their capability to sense reliably molecular interactions, we performed adhesion force spectroscopy experiments during two consecutive days (day 1 and day 2) on cells from the same passage plated on two different petri dishes. We observed that the probe detected similar mean number of jumps N_j_ (Figure S4A), with some discrepancies at higher contact times, and very similar force per jump <F_j_> (Figure S4B). Remarkably, besides the satisfactory agreement of the mean N_j_ and <F_j_> values, the distributions of the measured single-bonds forces for the two days for the same contact times agree very well (Figure 4A,B), and also clearly show that there is an increase of the force at higher contact times. This leads us to conclude that the same probe in different days is able to measure the same adhesive events, i.e. the probe maintains its ability to capture both qualitatively and quantitatively the biological picture of the adhesive cell-ECM interaction. The slight difference observed from day to day in the measured parameters could be explained by the well-known heterogeneity of cells, as well as by their high sensitivity to slight changes in the environmental conditions.

**Figure 4.**
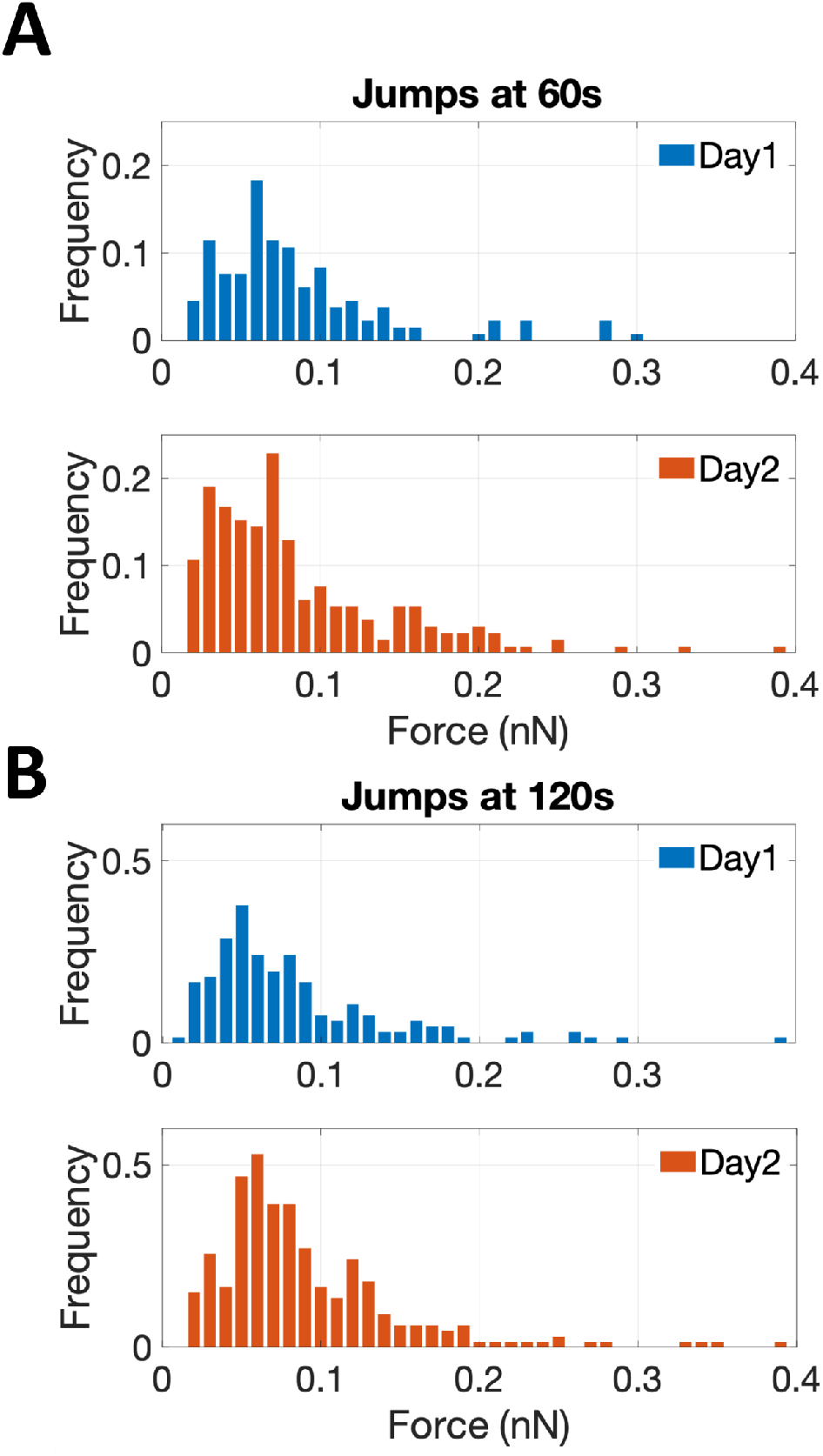
Results of the adhesion force spectroscopy measurements performed with the same ECM probe in two consecutive days (day 1 in blue and day 2 in red) on AY-27 cells from the same passage. The distributions of the force of single jumps at two contact times are shown in (A) (ct=60s) and (B) (ct=120s).

### 3.3 Use of tailored native ECM probes in adhesion force spectroscopy experiments

To demonstrate the potential of native ECM-probes for the study of cell-microenvironment interactions, we carried out adhesion force spectroscopy experiments with these probes against AY-27 cells.

For the proof-of-principle presented here, we selected two main bladder regions for the creation of the ECM probes: the submucosal region, closer to the lumen of the bladder, rich in connective tissue to support the attachment of the urothelium, a type of stratified epithelium made of transitional epithelial cells, and the detrusor region, a central layer of the bladder made of muscle fibres that allow contraction ^40,41^, as shown in Figure 5A-C. The ECM probes, named after their corresponding region of origin (Probe SM, submucosal, and Probe D, detrusor) are interesting because, albeit deriving from adjacent areas of the same bladder tissue, they are characterised by major differences in the composition and mechanical properties ^17,40^.

**Figure 5.**
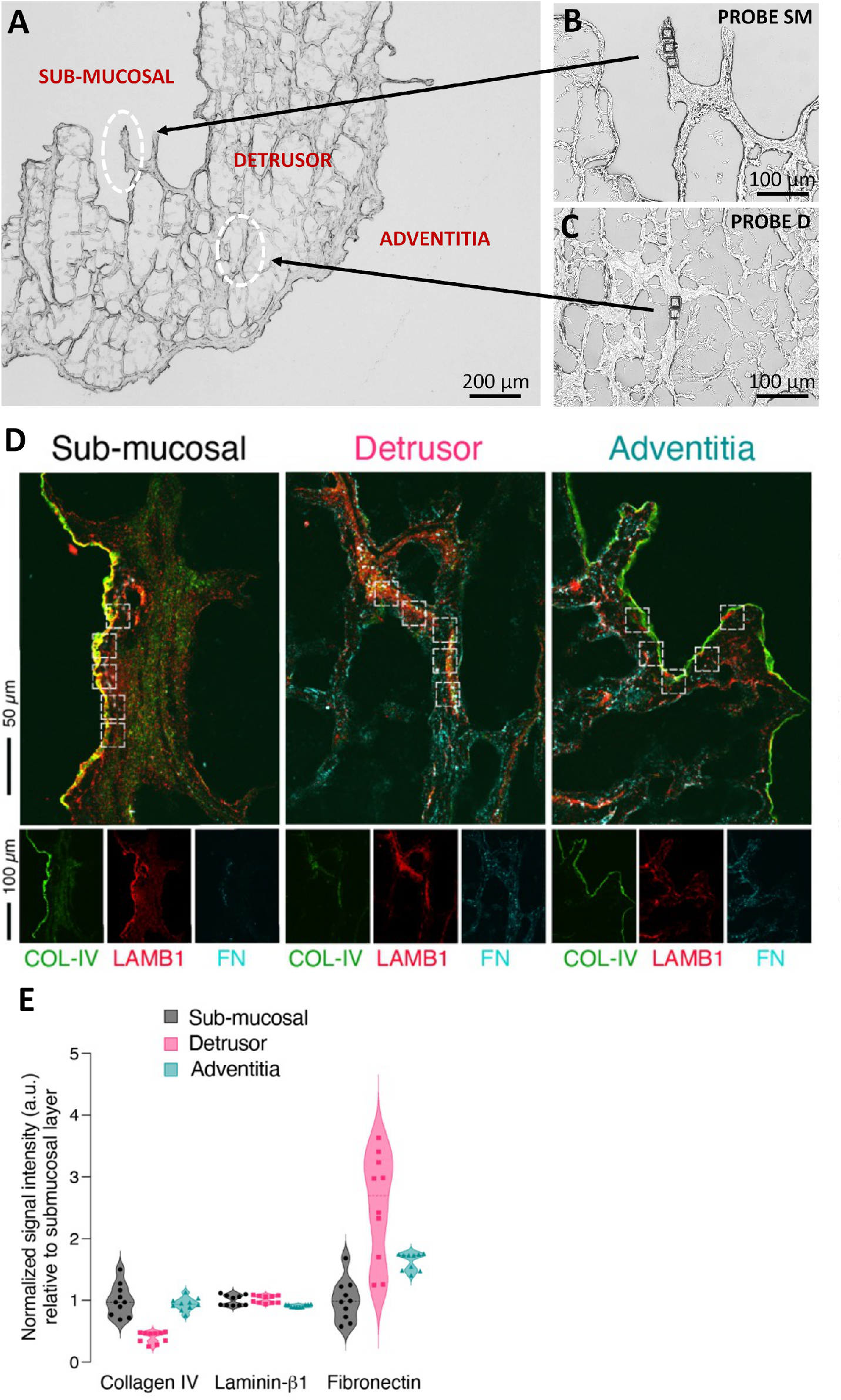
Optical images of the ECM before (A) and after (B and C) the LMD cut. The white circles highlight the regions of the cuts, from where the ECM probes have been produced: probe SM from the submucosal, and probe D from the detrusor. The denomination of the specific regions of the bladder in (A) follows Refs ^40,41^. (D) Immunostaining of different regions of the decellularised bladder (submucosal, detrusor and adventitia) with collagen IV (COL-IV, *green*), laminin β1 (LAMB1, *red*), and fibronectin (FN, *cyan*). (E) Quantification of the fluorescent signal associated to the different proteins in each ROI (white squares). The data represent normalised fluorescent signal (a.u., arbitrary units) relative to the mean value of the submucosal layer for each matrix (n=10, 5 ROIs per each field, 2 field acquired per each layer).

We performed immunostaining and quantification of the abundance of principal ECM proteins, such as collagen, laminin, and fibronectin, which are responsible for major integrin-mediated cell/ECM interaction in the bladder tissue (Figure 5D,E). We observed a lower frequency of occurrence of collagen IV in detrusor, compared to submucosal and adventitia regions of the ECM (Figure 5E).

While laminin seems to be consistent in all the matrix, fibronectin is more present in the detrusor compared to the other two proteins. We furthermore observed different mechanical properties of the ECM regions (Figure S5), in agreement with the results recently reported for the three tissue layers of the rat bladder (i.e., urothelium, lamina propria and muscle layer)^17^. These assessments confirm that the different parts of the bladder ECM reproduced by Probe SM and Probe D exhibit significant distinctions in their compositional and biophysical features.

We then performed adhesion force spectroscopy experiments comparing Probe SM and Probe D against AY-27 cells. For both substrates, N_j_ rose initially (0-60s) and then remained nearly constant. While no clear difference is seen in the number of jumps (Figure 6A), we observed important differences between the two ECM probes in both <F_j_> (Figure 6B) and F_a_ (Figure 6C). For both parameters, the values were higher for the SM substrate at almost all time points. The maximum adhesion force for Probe SM reached ~1.5 nN, compared to only ~0,5 nN for Probe D. A similar outcome was detected for <F_j_>.

**Figure 6.**
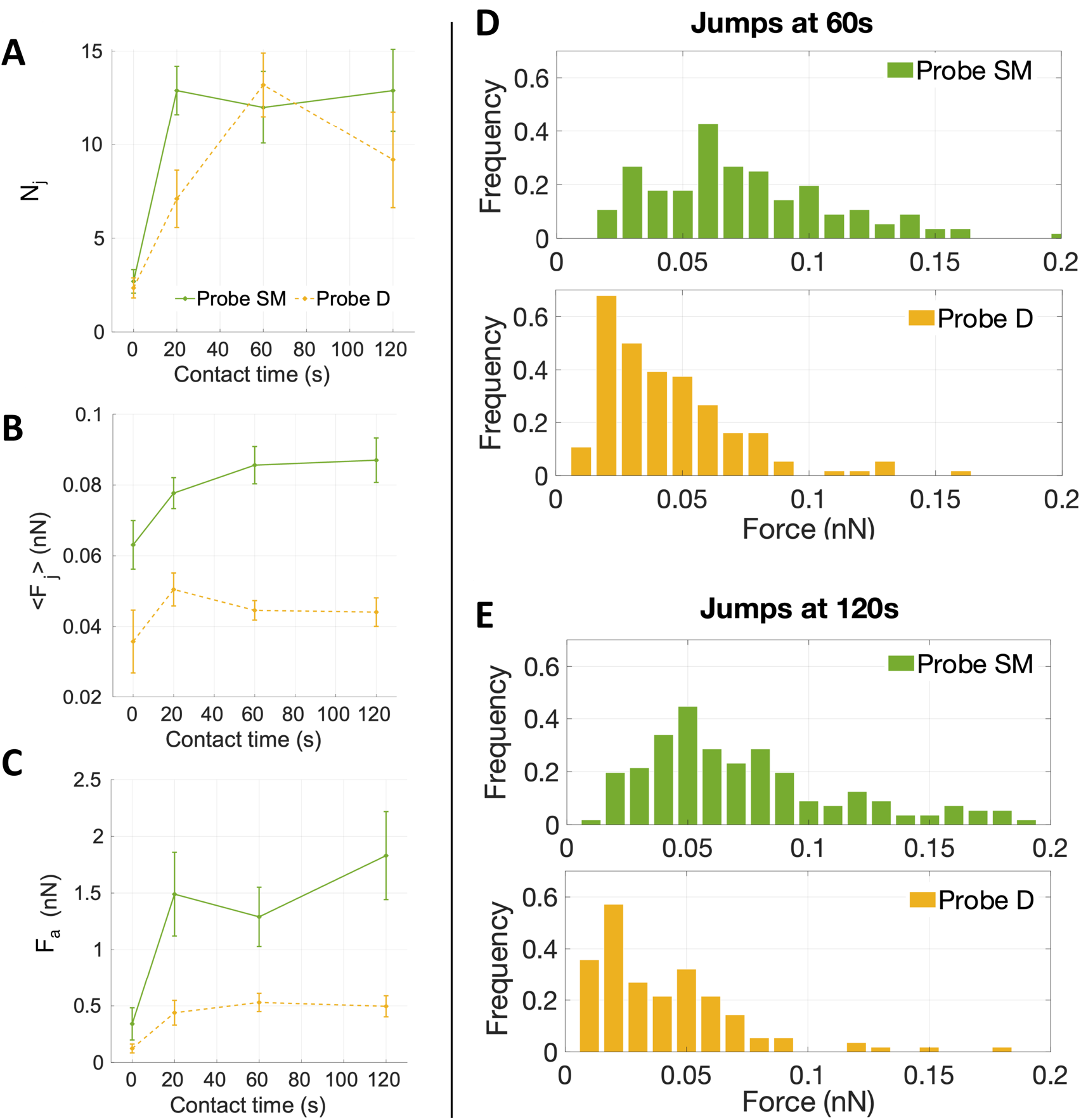
The figure shows the results of adhesion force spectroscopy measurements performed with two different ECM probes on AY-27 cells of the same passage: Probe SM (green) and D (yellow), obtained using ECM fragments from submucosal and detrusor regions, respectively. Four different cell/probe contact times have been tested (ct = 0 s, 20 s, 60 s, 120 s). (A-C) The mean number of events (jumps) N_j_, the mean force per jump <F_j_>, and the mean total adhesion force F_a_ as a function of the contact time, respectively. (D,E) The distributions of the force of single jumps at two contact times (ct=60s and 120s, respectively). Error bars in A-C represent effective standard deviations of the mean, as explained in the Methods.

Remarkably, the distribution of the forces of jump unbinding events (Figure 6D,E) are very different for the two probes. For Probe SM, a more ample spectrum with occurrence of higher forces per jump was noted, whereas for Probe D the forces are restricted to lower values. Moreover, at longer contact times we observed a moderate shift towards lower forces for Probe D, while in the distribution of Probe SM the frequency of both smaller and higher forces increased slightly.

These results demonstrate that our approach allows to produce ECM-probes from different regions of a tissue with tailored adhesive properties, to sense qualitatively and quantitatively different molecular interactions. The unbinding events might be attributable to different types of bonds, i.e., cell-fibronectin or cell-collagen IV interaction, depending on the ECM portion attached to the probe. Since AY-27 rat bladder tumour cells are derived from a urothelium transitional carcinoma,^44^ they likely express those receptors that bind ECM components present in the submucosal layer, since this region of the bladder allows urothelial cells attachment, as mentioned above. The probe derived from the fibronectin-dominated, muscular type detrusor layer is probably less ideal for AY-27 attachment therefore explaining the poorer affinity observed. The increase of the frequency at higher forces at longer contact times for Probe SM is congruent with the transition from single to cooperative α2β1 integrin receptor binding to collagen I during the early steps of adhesion, shown by Taubenberger et al^45^. In the fibronectin-prevalent Probe D condition this does not occur (the trend is more in the opposite direction). However, the forces measured for the first pristine adhesion events to Probe D at 0s are in the range of what was measured for α2β1 integrin/fibronectin binding by Sun et al. (39±8 pN versus 36±9 pN in our measurements, see Table 1). The lower binding force and the lower adhesion pattern measured with Probe D are consistent with the invading character of cancer cells in the inner muscular detrusor layers of the bladder^46,47^, which is necessary for the tumour to become muscle invasive. Altogether, the specific adhesive features measured with the two ECM probes are in line with the composition of the native ECM portions (Figure 5D,E) and distinct types of cell-ECM interactions.

**Table 1.**
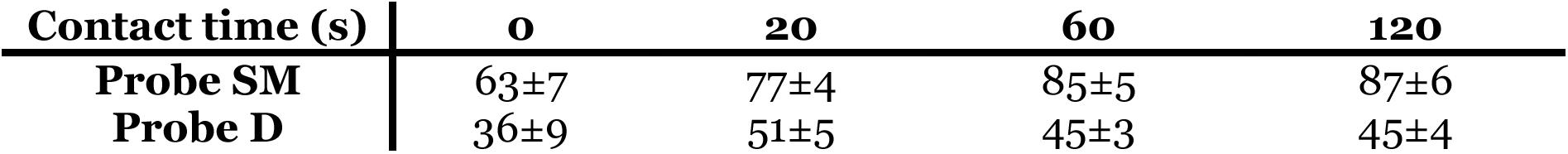
Mean forces per jump at contact times 0, 20, 60 and 120 seconds.

| Contact time (s) | 0 | 20 | 60 | 120 |
| --- | --- | --- | --- | --- |
| Probe SM | 63±7 | 77±4 | 85±5 | 87±6 |
| Probe D | 36±9 | 51±5 | 45±3 | 45±4 |

To validate the association of measured jumps to integrin-related events, we performed a control experiment treating the cells with an allosteric inhibitory antibody against β1 integrin (4b4), at fixed contact time (ct=60 s), using two ECM probes made from a submucosal region (Probe CTRL1 and CTRL2, Figure S6). The major component of the submucosa is collagen IV (Figure 5D,E), which is bound by α1β1 and α2β1 integrins, targeting β1 was then strategical. We observed force of adhesion, mean force of jumps and tethers and mean number of events before and after the addition of 4b4 (Figure 7). We considered both jumps and tethers as representative events in these experiments, since the inhibitory antibody targets all β1 subunits of the integrins, either linked to the cell cytoskeleton (jumps related events) or not (tethers)^45,48–50^. The two ECM-probes CTRL1 and CTRL2 (Figure S6) were used on two consecutive days and data were grouped together, as it was observed that the force of events were similar and displayed similar distributions (Figure S7).

**Figure 7.**
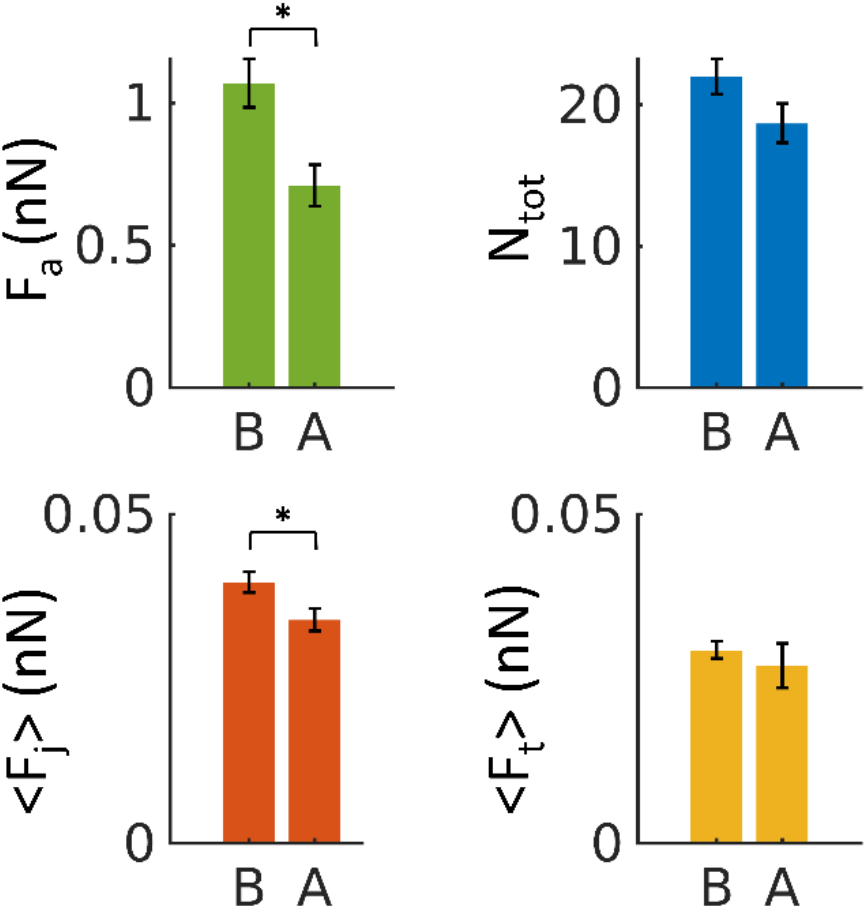
The mean force of adhesion F_a_, mean number of events N_tot_ (both jumps and tethers) and mean force of jumps and tethers measured <F_j_> and <F_t_> respectively, using control probes CTRL1 and CTRL2 before (B) and after (A) the addition of the 4b4 inhibitory antibody against β1 integrin at concentration of 5μg/ml. Errors represent effective standard deviations of the mean, as explained in the Methods.

Figure 7 shows a significant decrease upon exposure of cells to the antibody of the total adhesion force F_a_ as well as the mean force of jumps <F_j_>. A decrease of the total number of events N_tot_ (jumps and tethers) and the mean force of tethers <F_t_> was also observed, although not significant. These results confirm that the measured interaction forces depend at least partially on β1 subunits of integrin proteins.

## 3. Conclusions

In the current work, we presented a novel adhesion force spectroscopy approach for the study of the cell-microenvironment interaction obtained by attaching selected pieces of native ECM to an AFM tipless cantilever, exploiting laser microdissection. These probes reproduce the full complexity of the cell-ECM interface in the physiological condition, in terms of both biophysical cues and composition.

We demonstrated, as proof-of-principle, the functionalisation of AFM cantilevers with native rat bladder ECM pieces from different regions of the tissue, and the use of these novel probes in adhesion force spectroscopy experiments against bladder cancer cells AY-27.

With this novel strategy, we could highlight differences in the adhesive pattern between the AY-27 cells and the ECM when isolated from the submucosa or the detrusor layer. The lower adhesion force detected when the detrusor ECM was used highlights the relevance of our strategy to investigate the impact of cell-ECM adhesion on tumour invasion ^46,47^. An added value of our approach is that its spatial resolution allows measurements at the single cell level, and thus it could be used to assess the heterogeneity of neoplastic clones.

The robustness and durability of the ECM probes were demonstrated by stress tests performed both in ramping and contact scanning modes. In addition, we showed the ability of this approach to reliably and repeatedly detect specific integrin-related events, as well as to assess significant differences in the mechanotransductive parameters such as the mean number and force of unbinding events, and their distribution, depending on the ECM region used to produce the probe. The observed differences correlate with the differences in the ECM protein and collagen composition and biophysical properties, revealed by the immunostaining and mechanical analysis.

Our results demonstrate the potential of these novel AFM probes to perform highly specific force spectroscopy investigations. Indeed, the adhesive properties of the probes can be tailored by selecting the type and the specific region of the ECM. The target cells can also be selected to probe specific mechanotransductive and mechanobiological interactions.

For example, experiments can be performed probing different types of matrices against a wide variety of cells (both healthy, tumoural, and from metastatic sites). Both ECM and cells could be extracted from different organs and tissues of the same patient, which might find application in identifying biomarkers for early diagnosis of diseases (particularly the context of cancer and metastasis, but also other diseases). This could allow in-depth understanding of disparities in the cell/microenvironment interaction between healthy, tumoural and metastatic cells confronted with different ECM substrates. The approach can also be used to test the effects of chemotherapeutic drugs targeting mechanotransduction-associated key players (such as integrins or Rho signalling-related proteins) and structures (e.g., components of the ECM, glycocalyx, and cytoskeleton) in pathophysiological conditions, with a very high precision at the level of forces and number of adhesion sites.

This novel adhesion force spectroscopy approach is therefore characterised by a high potential versatility, which could be a key element in the nanotechnological toolkit of nascent personalised medicine.

## 4. Material and methods

### Cells

The rat bladder cancer cell line AY-27 (Sigma-Aldrich, cat. number SCC254) was authenticated for lack of cross-contamination by analyzing 9 short tandem repeats DNA (IDEXX Bioanalytics, Ludwigsburg, Germany)^51^. The cell line was cultured in RPMI 2mM L-glutamine with 10 % FBS, 1% penicillin/streptomycin and 1% amphotericin; cells were cultured in an incubator at 37°C and 5% CO_2_ (Galaxy S, RS Biotech). All reagents and materials are from Sigma Aldrich, if not stated otherwise.

For AFM measurements, the cells were plated one day before on glass bottom petri dishes (∅ 40 mm Willco Wells) coated with poly-l-lysine (0.1% w/v for 30 min at room temperature), in the same RPMI medium without phenol red.

### Extracellular Matrices

The bladder from healthy rat was decellularised following the protocol from Genovese et. al ^52^, validated also for bladder^53^. The ECM samples were embedded in optimal cutting temperature compound (OCT), frozen in dry ice and kept at −80°C. All procedures and studies involving mice were approved by the Institutional Animal Care and Use Committee of San Raffaele Scientific Institute, and performed according to the prescribed guidelines (IACUC, approval number 942). Adult (9-10 weeks old) female Fischer rats were from Charles River Laboratories, Italy.

For the laser microdissected slices, 10μm thick ECMs were cut with a cryostat and mounted onto classical optical microscopy glass slides. The sample was then immerged in PBS to remove OCT and then in Ethanol 70% to dehydrate it prior to laser microdissection.

We tested the effect of dehydration of the ECM and we confirmed that this process does not alter the mechanical properties of the matrix. For the comparison of mechanical properties before and after dehydration, an ECM slice was first characterised after gently washing away the OCT with PBS, then dehydrated in 70% EtOH (following the same procedure used for LMD) and measured again (Figure S3C).

Two different slices were used for the comparison of morphological and mechanical properties of the inner and outer regions of the LMD cut.

For AFM mechanical experiments, 50μm thick ECMs were cut with a cryostat and attached to super-frost microscope glass slides. The prepared ECM slices were kept at −20°C prior to the AFM experiments. Before starting the AFM measurements, the slices were washed with PBS to remove the OCT.

### Reagents

For the functionalisation of the probes, 3-aminopropyl triethoxysilane (APTES) and Genipin were used, both purchased from Sigma. Genipin was diluted in 1,5% (v/v) in PBS. 4b4 antibody was purchased from Beckman Coulter.

### Laser Microdissection (LMD)

400 μm^2^ (20μm x 20μm) regions were cut using a UV-based LMD7 laser microdissection system (Leica Microsystems) at 20x magnification placing the slide with the ECM section facing up. This setup allowed to separate the regions of interest (ROIs) from the rest of the tissue without detaching them from the glass. The engraved sections were kept dry at 4°C until further processing.

### Atomic Force Microscopy

All the experiments have been performed using a Bioscope Catalyst AFM (Bruker), mounted on top of an inverted optical microscope (Olympus X71). The system was isolated from the ambient noise placing the AFM on top of an active antivibration base (DVIA-T45, Daeil Systems) and enclosing it in an acoustic box (Schaefer, Italy).

#### Mechanical measurements

aimed to characterise the Young’s modulus of elasticity of the ECM were performed by recording force versus distance curves (shortly force curves, FCs), then transformed into force vs indentation curves, as described in Refs ^13,54,55^. We used custom colloidal probes, produced by attaching borosilicate glass spheres to tipless cantilevers (MikroMasch HQ:CSC38/Tipless/No Al or NanoandMore TL-FM), as described in Ref. ^56^. The tip radius was calibrated by means of reverse AFM imaging ^56^. The cantilever spring constant was calibrated using the thermal noise method ^57,58^, and fine corrections were applied to account for geometrical and dimensional issues ^59,60^. The deflection sensitivity (or inverse optical lever sensitivity, invOLS) of the optical beam deflection apparatus was measured as the inverse of the slope of the deflection vs z-piezo displacement curves acquired on a stiff substrate ^55^..The deflection sensitivity was monitored and if necessary corrected during AFM experiments using the contactless SNAP method ^61^, assuming as reference spring constant the intrinsic spring constant previously calibrated.

The probes used had spring constants of 3-4 N/m, radii of 5.5 and 9 μm. 10 sets of force-distance curves (force volumes, FVs), consisting of 20×20 curves each, spanning typically an area of 50μm x 50μm, were acquired on regions separated by more than 200 μm on each slice. All measurements were carried out in a droplet of PBS, confined on the glass slide by means of hydrophobic ink (Sigma).

The precise alignment of AFM and optical images was possible thanks to the Bruker MIRO software, which allowed to choose the ROIs on the ECMs for the analysis (as during the force spectroscopy experiments). The transparency of the slices with thickness below 50 μm allowed us to identify and select regions where pieces of ECM had been cut by LMD. The ECM probe can be placed precisely on top of a cell using the optical alignment, ensuring that contact is established with the ECM, and not with another part of the cantilever.

#### Adhesion Force Spectroscopy measurements

were performed according to the protocol described in Refs ^11,25^, by acquiring several FCs per cell on at least 4 cells per each contact time, using the ECM probes. In force spectroscopy experiments, the adhesion pattern in the retracting portion of the FC is studied; upon contact with the sample, the tip was retracted after a contact time (ct) of 0, 20, 60 and 120s, during which the z-sensor position was kept constant by the feedback of the AFM. A typical retraction portion of a FC is shown in Figure S8, where the adhesive features (jumps and tethers) can be clearly observed. We typically used the following values for the ramp parameters: ramp frequency, 1 Hz; ramp size, 10 μm; maximum force, 1nN.

For the control (inhibition) experiment, the 4b4 antibody, an inhibitor of β1 integrin, was added to the medium, at a concentration of 5 μg/mL, 20 minutes before the AFM measurement. Experiments were carried out at 37°C using a thermostatic fluid cell and a temperature controller (Lakeshore 331, Ohio, USA).

#### Functionalisation of tipless cantilever

The tipless cantilevers were cleaned in an oxygen plasma chamber at a power of 80W for two minutes prior to the functionalisation, to remove organic contaminants and maximise the number of surface -OH groups. The vapor APTES deposition was performed under static nitrogen for three minutes in a desiccator with 50μl of APTES (Sigma). Cantilevers are then washed in toluene to remove any unbound APTES and left in an oven for the curing of the functionalisation ^62,63^. The successful functionalisation of the cantilevers with APTES was checked by wettability measurements on an equivalent silicon substrate (with native oxide layer on top), before and after APTES deposition (Figure S9). This substrate has the same surface chemistry of AFM cantilevers. Deposition of APTES typically makes a hydrophilic surface more hydrophobic, as revealed by the marked increase of the contact angle in Figure S9. On APTES functionalised cantilevers, we deposited a droplet of genipin for 20 min to allow covalent reaction both with APTES and future ECM ^64,65^. Afterwards, the cantilevers were gently washed with PBS and directly used for assembling with the ECM pieces. The obtained probes can be reused many times; indeed, the removal of the functionalisation can be done by piranha cleaning, as discussed in Supporting Note SN1^66^.

#### Production of native ECM probes

The laser microdissected pieces of ECM (Figure S2 and S6) were attached to functionalised tipless cantilevers (MikroMasch HQ:CSC38/Tipless/No Al). Attachment of ECM pieces to tipless cantilevers, rather than to cantilevers with tip, was found to be more reliable. The procedure for the detachment of ECM pieces from the glass slide and the attachment to the tipless cantilever is described in detail in the Results section. For force spectroscopy experiments, the spring constant of the cantilevers was chosen in the range 0.01-0.05 N/m.

#### Data analysis

Processing of the data was carried out using custom routines written in Matlab (Mathworks) language. The raw FCs, consisting of the raw deflection signal from the photodetector (in Volts) units as a function of the z-piezo displacement (in nm), have been rescaled using the measured calibration factors (deflection sensitivity and spring constant) into force (in nN) vs indentation or tip-sample distance (in nm), according to the standard procedure ^55^.

The elastic properties of cells and ECMs were characterised through their Young’s modulus (YM) of elasticity, extracted by fitting the Hertz model to the 20%-80% indentation range of the FCs (details in Refs ^13,54^),

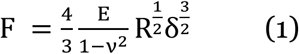
which is accurate as long as the indentation δ is small compared to the radius R. In Eq. (1), v is the Poisson’s coefficient, which is typically assumed to be equal to 0.5 for in-compressible materials, and E is the YM.

Finite thickness correction for cells was applied as described in Refs ^54,67,68^. In the case of ECM, despite the relatively small thickness of the slices, this correction was not applied because the thickness is not known with good accuracy, being the underlying substrate not always accessible; nevertheless, despite a systematic overestimation of the YM of the ECM, we were able to carry out a comparative characterisation of ECM elasticity in different conditions.

For the analysis of adhesion force spectroscopy data, a custom MATLAB routine was used to detect specific adhesion events in the FCs (jumps and tethers ^22,25^) and to calculate the values of the relevant parameters (see Figure S8), as described in Chighizola et. al. ^25,69^, like the mean number of jumps and tethers per force curve, N_j_ and N_t_, respectively, the total number of events N_tot_=N_j_+N_t_, and the mean total adhesion force F_a_, all these quantities being averaged across all FCs for a given condition; the mean force per jump <F_j_> and the mean force per tether <F_t_> represent the mean values from the distribution of F_j_ and F_t_ values measured in a specific condition. The associated errors were calculated by summing in quadrature the standard deviation of the mean to an instrumental error of 3%, calculated by propagating the calibration uncertainties in the fitting procedure through a Monte Carlo simulation^54^.

### Immunofluorescence

10μm thick snap-frozen OCT-embedded cryosections were mounted on positively charged glass slide (Superfrost plus adhesion slide, #J1800AMNZ, Epredia Inc) and processed as described above for laser microdissection. Engraved and non-engraved sections were fixed in 4% Paraformaldehyde (#15710 EM grade, Electron Microscopy Sciences) at room temperature for 10 min. After washing in PBS, sections were incubated for 1 hour with blocking buffer (2% Donkey serum, 1.5% BSA, 0.25% Fish Gelatin, PBS pH=7.2) and then incubated overnight at 4°C with the following primary antibodies: anti-Laminin β1 (4μg/ml, sc-17810, Santa Cruz Biotech), anti-Fibronectin (10 μg/ml, NBP1-91258SS, NovusBio), anti-Collagen IV (20 μg/ml, AB769, Millipore).

After washing, anti-goat Alexa Fluor® 488, anti-mouse Alexa Fluor® 647 and anti-rabbit Alexa Fluor 555 conjugated fluorophore-labeled F(ab)2 donkey secondary antibodies were used (Jackson Immunoresearch). Sections were DAPI counterstained and 3×3 Z-stacked large images (9 mm depth, 11 stacks) were acquired using a Yokogawa Spinning Disk Field Scanning Confocal System (CSU-W1, Nikon Europe BV, Amsterdam, Netherlands) equipped with 405, 488, 561, 640, 785 nm lines of solidstate lasers, 40x/1.15NA water immersion objective lens and a Prime BSI sCMOS camera (Teledyne Photometrics, Tucson, AZ).

Mean fluorescent intensity signal was quantified in multiple ROIs (175 um2 each; n=10, 5 ROIs per each field, 2 field acquired per each layer) and normalised to the respective average pixel intensity of the entire stained area. Data represent the normalised signal intensity compared to the average value of the submucosal layer for each matrix. For pre- and post-cut evaluation, fluorescent signal intensity was quantified in the central portion of the engraved region (Post-LMD) and the corresponding area on a consecutive section not subjected to laser cut (Pre-LMD). The normalised signal intensity was averaged for each ECM and scaled between 0 and 1 (n=30, 5 ROIs per each field, 6 fields acquired distributed along the layers of the bladder). Mann-Whitney test was used to compare differences in the signal intensity.

## Author contributions

Conceptualisation: AP, CS;

methodology – ECM preparation: IL;

methodology – probe fabrication and characterisation: HH, GRD, AP;

methodology – laser microdissection and staining: LN, PDC, GRD;

methodology – cell culture and preparation: HH, CS;

methodology – AFM and force spectroscopy: HH, MC, AP;

data curation and analysis: HH, MC, GRD, PDC, AP;

original draft writing and editing: HH, MC, AP;

draft revision: all authors;

supervision: MC, GRD, CS, AP;

resources, funding, and project administration: AP, GRD, MA.

Author contributions were allocated adopting the terminology of CRediT Contributor Roles Taxonomy.

## Conflicts of interest

There are no conflicts to declare.

## Acknowledgements

This research was funded by the European Union’s Horizon 2020 research and innovation program under the Marie Skłodowska-Curie Action grant agreement No. 812772, project Phys2Biomed, and under FET Open grant agreement No. 801126, project EDIT.

PDC is supported by an AIRC fellowship.

## Supporting Information

**Figure S1.**
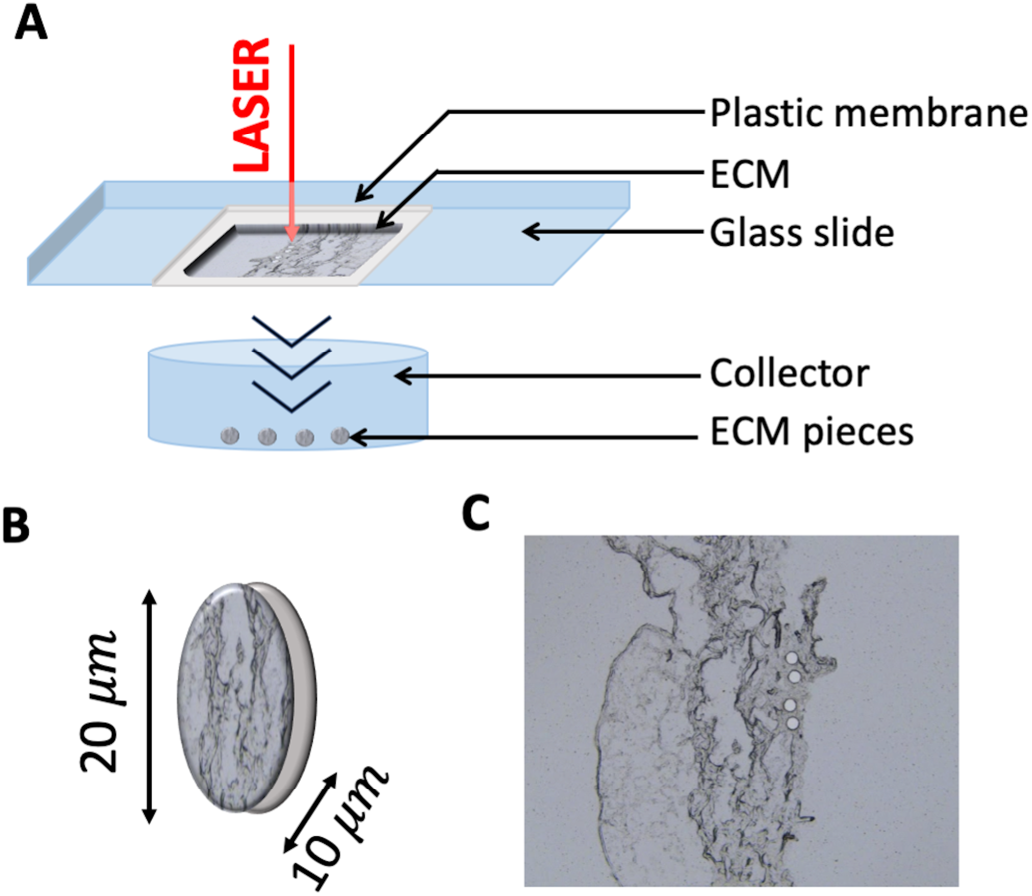
Schematics of the conventional use of the LMD system. (A) ECM fragments are cut by the UV laser together with the underlying plastic membrane and collected in a suitable reservoir where they fall by gravity. (B) The resulting circular pieces are 20 μm in diameter and 10μm thick, excluding the plastic membrane. (C) Upon cutting and collecting the pieces, circular holes are left in the ECM slice.

**Figure S2.**
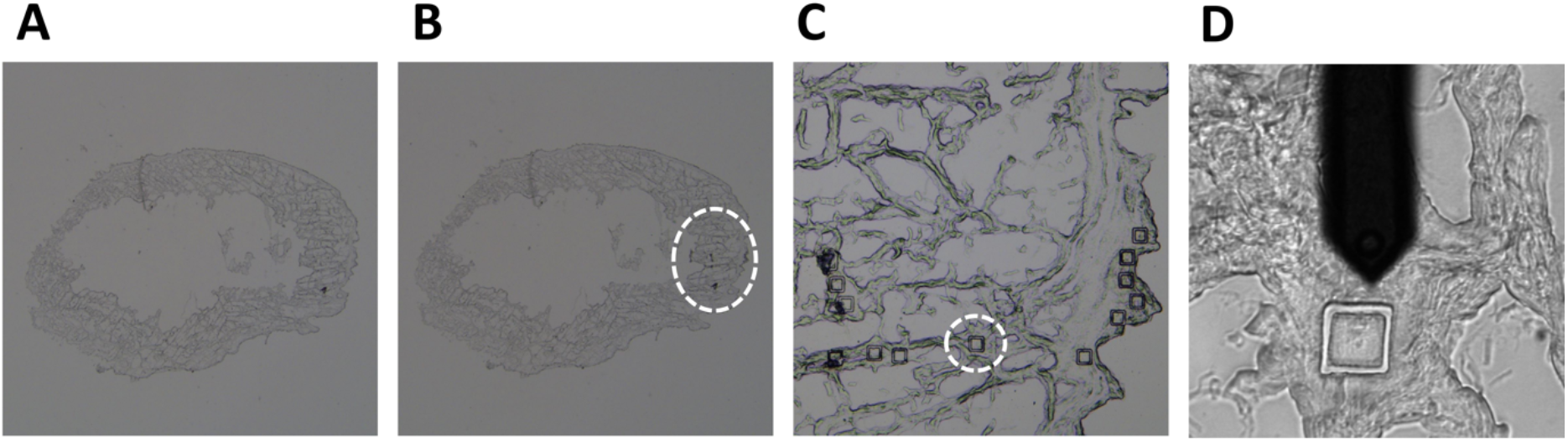
Optical images of the ECM rat bladder. (A) Before LMD cut, after PBS wash and ethanol dehydration. (B) After the LMD cut; the region of the cut is indicated. (C,D) Magnification of the cut region. The AFM cantilever is visible in D.

**Figure S3.**
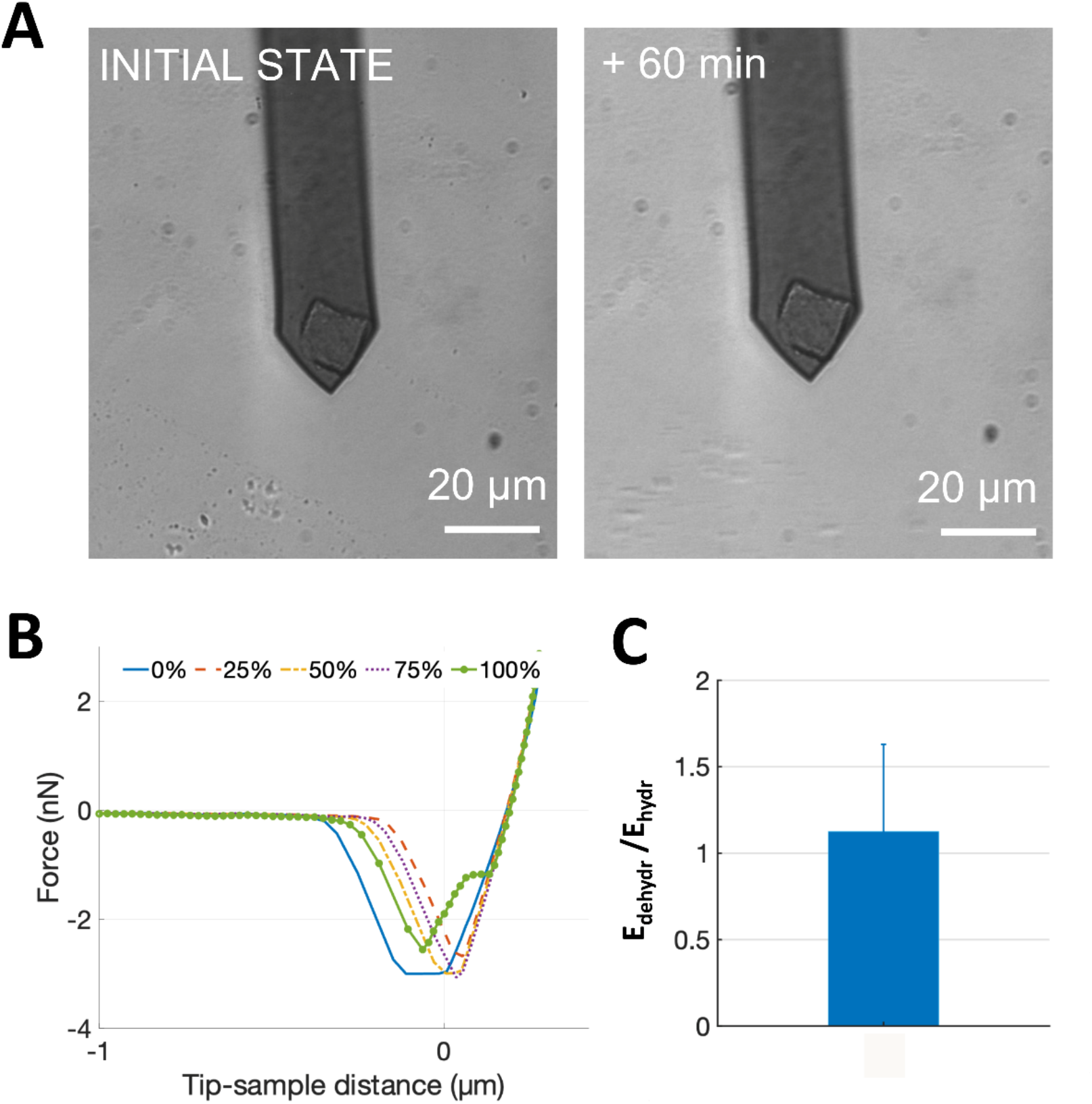
Stress tests on the ECM probes. (A) Optical images of an ECM probe before and after continuous scanning across a 20μm x 20μm area in Contact Mode for 60 minutes on a glass surface with a scan frequency of 1Hz. (B) Representative retraction curves acquired using an ECM probe ramped against a glass substrate before (0%), during (25, 50 and 75%), and at the end (100%) of an adhesion force spectroscopy experiment consisting in the acquisition of a total of 400 FCs. (C) The ratio E_dehydr_/E_hydr_ of Young’s modulus values E of an ECM sample before and after dehydration with 70% EtOH (the error bar represents the error calculated as the propagation of the effective standard deviations of the mean of E_hydr_ and E_dehydr_).

Stress tests were performed to assess the firm attachment of the matrix to the cantilever and the force spectroscopy functionality after repeated use.

Firstly, we perform continuous scanning of a 20μm x 20μm square on glass at a frequency of 1Hz with the probe for more than 60 minutes and observed no appreciable displacement or damage (Figure S3A). Then, we carried out force spectroscopy in liquid on a glass surface, with 10s contact time, by recording 400FCs at 1Hz with 5nN applied force. Looking at representative FCs at the beginning (0%), during (25, 50 and 75%) and at the end of the experiment (100%), we did not notice important changes in the measured total adhesion force (Figure S3B) (a slight decrease can be noted at the end of the test, while no significant differences is observable in the first 75% of the test). Interestingly, in this case, where the ECM is interacting with a glass surface instead of a cell, we did not detect unbinding events (jumps and tethers) in the FCs, which are typically associated to integrin-related specific adhesive interactions.

Furthermore, as LMD requires the dehydration of the matrix with a minimum of 70% Ethanol, we performed AFM nanoindentation measurements on the same ECM before and after dehydration with Ethanol. No significant difference in the Young’s modulus of the ECM was observed (Figure S3C).

**Figure S4.**
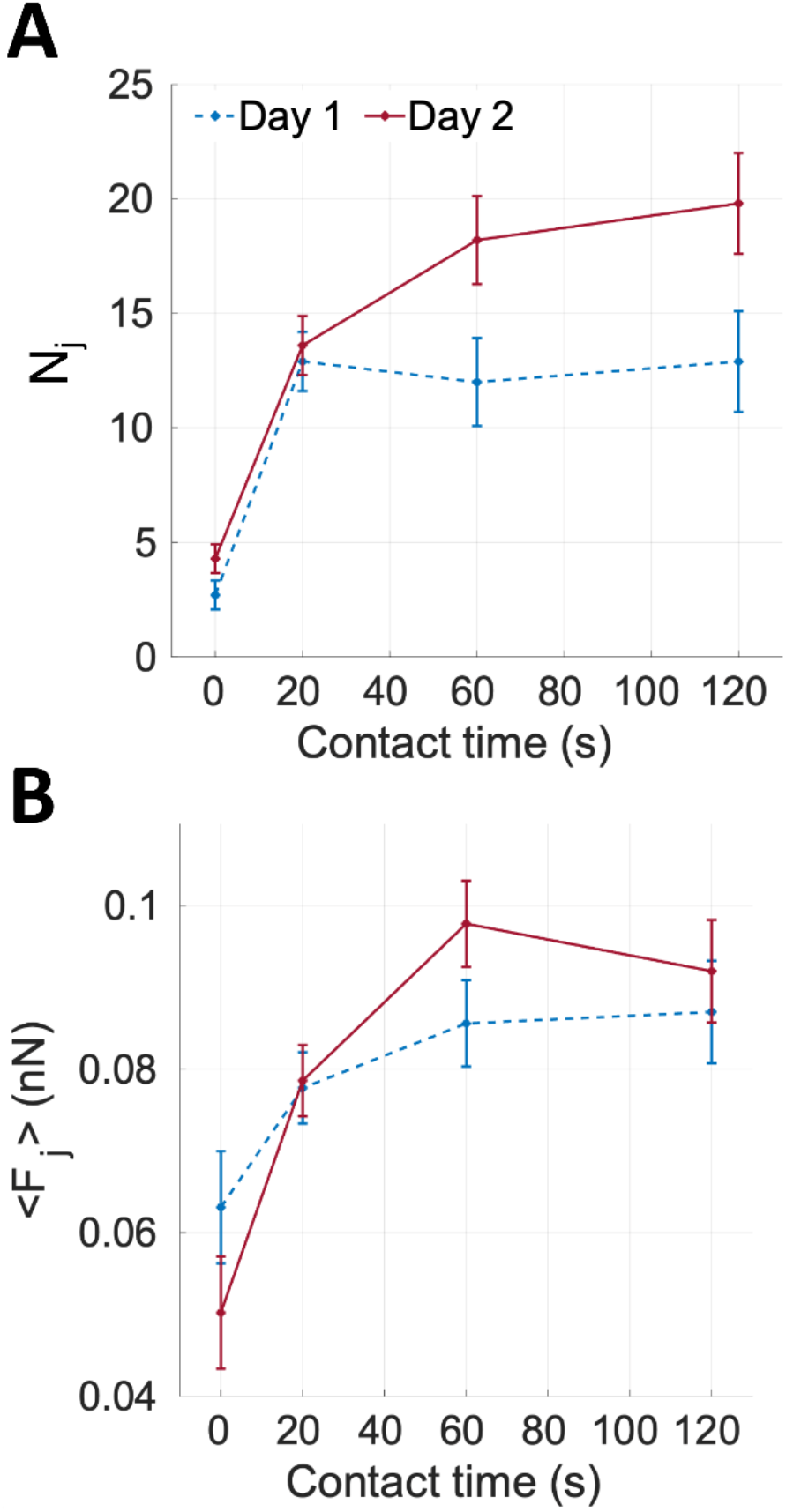
Mean number of jumps and force per jump measured on consecutive days. (boxes A and B, respectively), during adhesion force spectroscopy experiments performed with the same ECM probe on AY-27 cells from the same passage: day 1 (in blue) and day 2 (in red). Different contact times (ct) have been used (ct = 0 s, 20 s, 60 s, 120 s).

**Figure S5.**
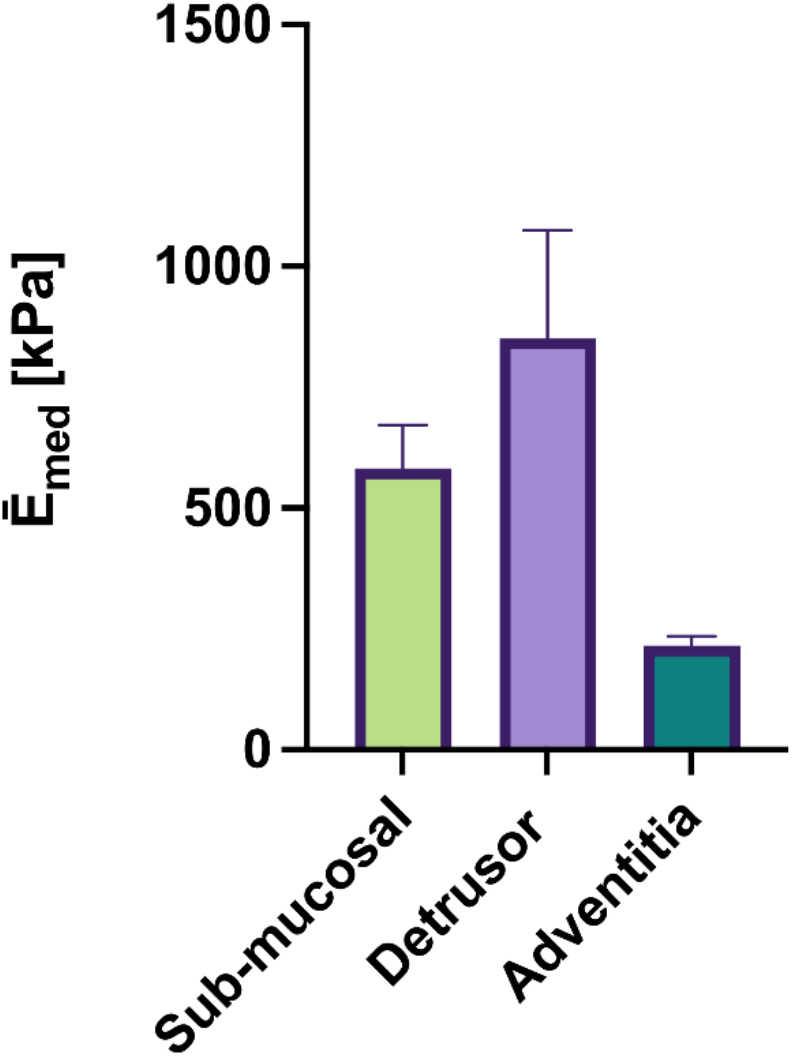
Young’s modulus of the different ECM regions: sub-mucosal, detrusor and adventitia. The mean median values are reported; error bars represent standard deviations of the mean. Data have not been corrected for bottom effect and might therefore overestimate the true Young’s modulus values. All measurements have been performed on the same 10μm decellularised ECM slice, on microdissected pieces (inside and outside taken as equivalent). Error bars represent the standard deviation of the mean.

**Figure S6.**
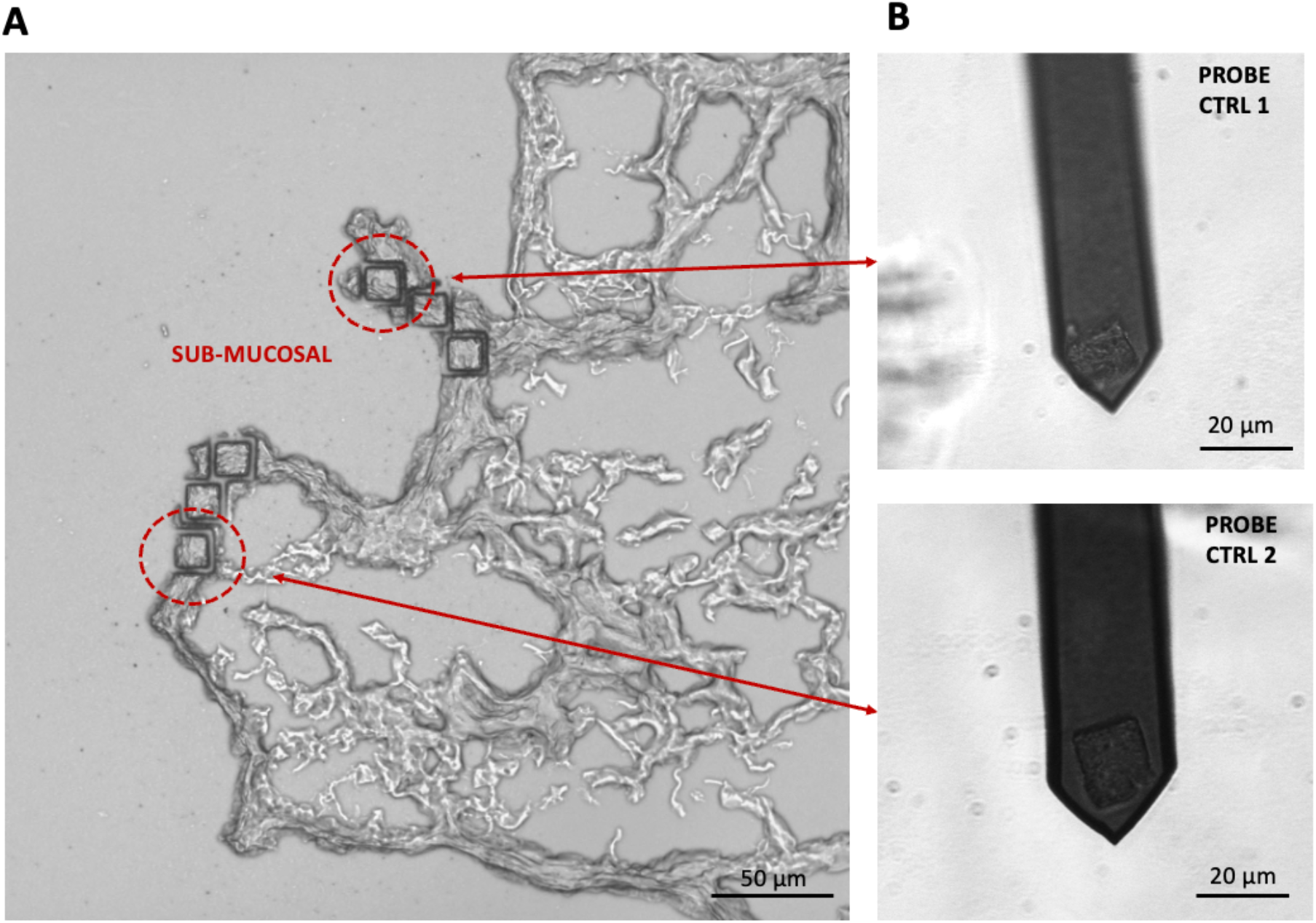
Optical image of the rat bladder ECM with the LMD cut. The red circles in (A) highlights the region in the submucosal layer where the ECM was cut to produce the control probes CTRL1 and CTRL2, shown in (B). The control probes were used in the integrin inhibitory experiments.

**Figure S7.**
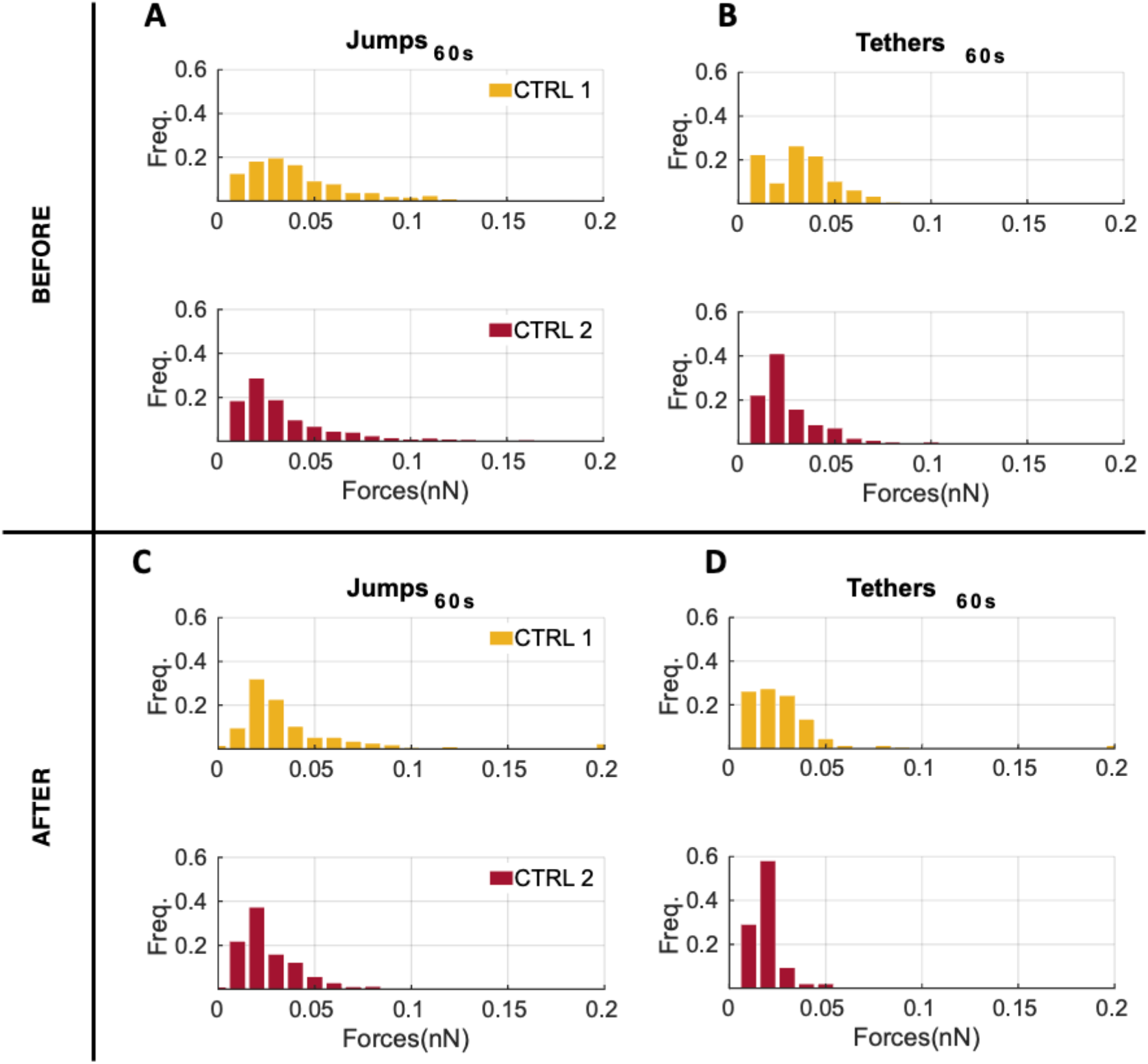
Distributions of jumps and tethers forces before and after the addition of the 4b4 integrin inhibitor antibody on AY-27 cells. (boxes A-B and C-D, respectively), measured during control adhesion force spectroscopy experiments with ct= 60s performed with the CTRL1 and CTRL2 probes (see Figure S6).

**Figure S8.**
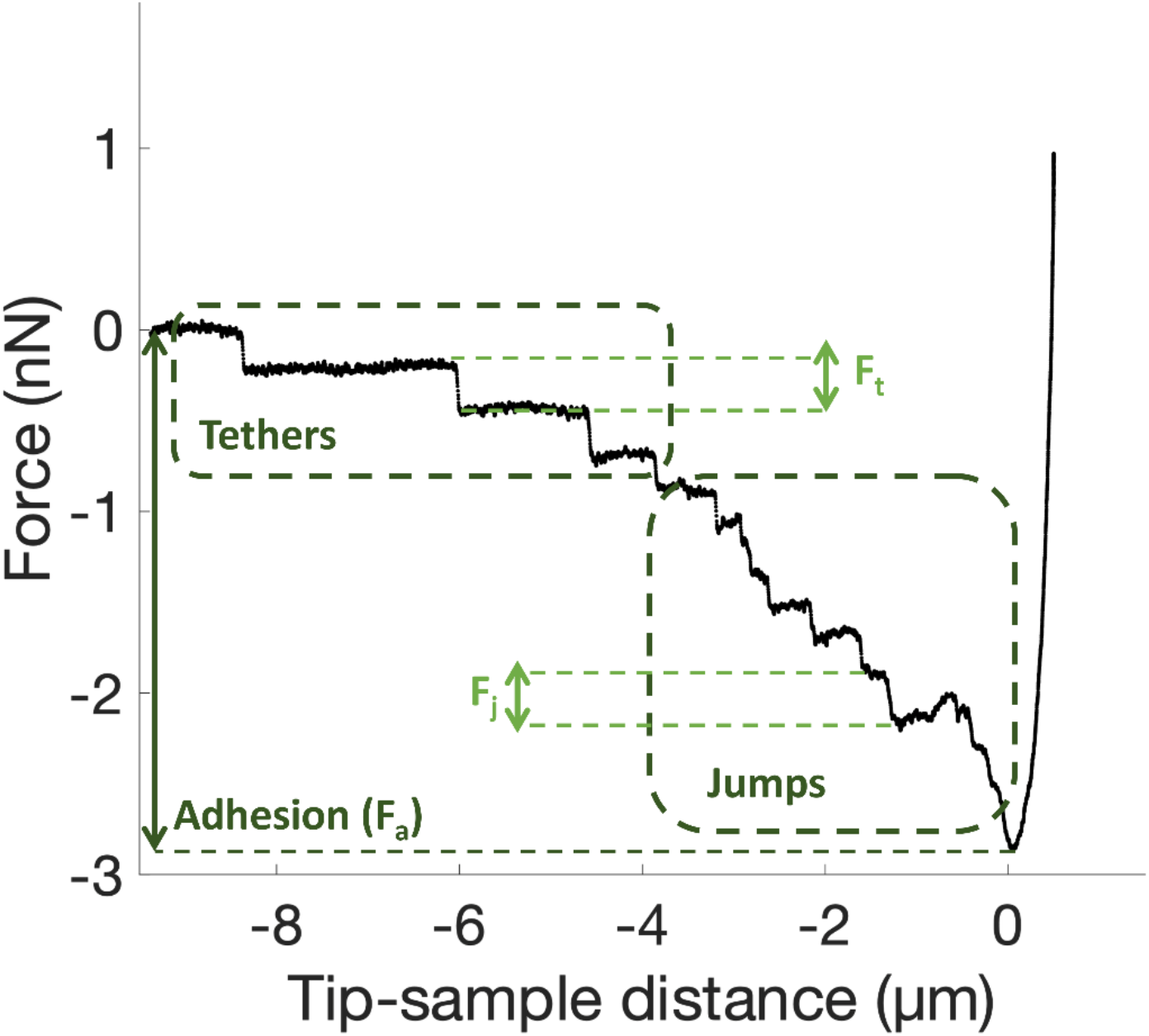
Representative force curve acquired during an adhesion force spectroscopy experiment using a native ECM probe. The relevant features and parameters are indicated: unbinding events (jumps and tethers), and the total adhesion force F_a_. The forces F_j_ and F_t_ necessary to break a single integrin-ECM bond in jumps and tethers, respectively, are shown. From the distribution of jumps and tethers forces, the mean jumps and tethers forces <F_j_> and <F_t_>, as well as the mean number of jump and tether events per force curves, N_j_ and N_t_, and the total number of integrin-related events N_tot_=N_j_+N_t_, are be calculated. The adhesion force F_a_ is calculated as the absolute difference between the minimum force value and the baseline value.

**Figure S9.**
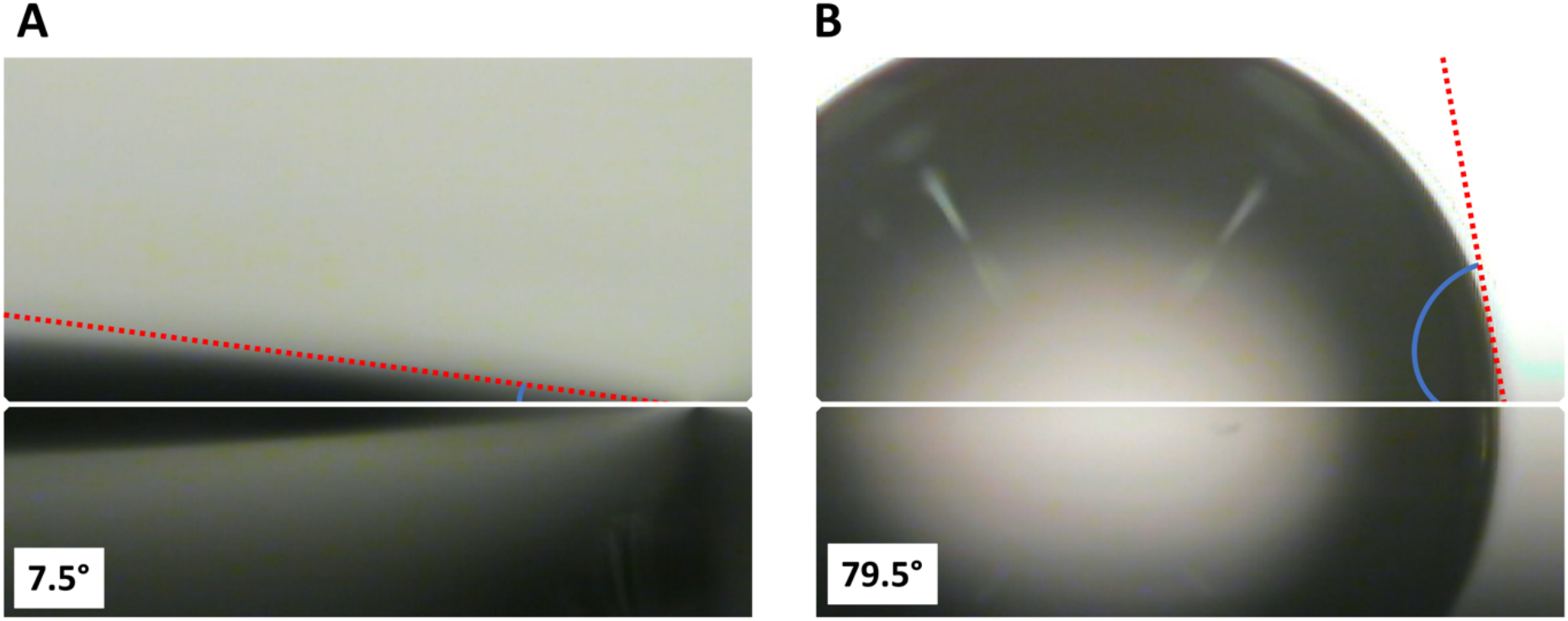
Contact angle analysis of water on functionalised silicon substrates, (A) before, and (B) after functionalisation with APTES. The contact angle increased from 7° to 79.5° upon APTES functionalisation of the oxidised silicon substrate.

**Supporting Note SN1. Cleaning and re-use of the ECM probes**

ECM probes can be cleaned and re-functionalised. In fact, the ECM can be degraded with proteolytic enzyme solutions such as helizyme, while the APTES coating (as well as the ECM) can be removed by piranha solution. Piranha solution is known to degrade any organic residues out of substrate such as silicon wafer and is commonly use in the AFM community for cleaning the probes.

**Supporting Movies SM1, SM2, SM3. Attachment of ECM pieces to functionalised cantilevers**

## Notes

### Competing Interest Statement

The authors have declared no competing interest.

### Summary of Updates

Updated Figure S5. Minor revisions (typos corrected).

